# APP and APLP2 Kunitz Domains Are Potent Endogenous Inhibitors of TMPRSS2 and Respiratory Virus Infection

**DOI:** 10.64898/2026.08.13.744607

**Authors:** Jan Lawrenz, Shreyans Chatterjee, Armando Rodríguez Alfonso, Annelies Stevaert, Rayhane Nchioua, Gina-Marie Templin, Giorgio Fois, Manfred Frick, Lieve Naesens, Rüdiger Groß, Jan Münch

**Author notes:** Corresponding author: Jan Münch, Institute of Molecular Virology, Meyerhofstrasse 1, 89081 Ulm, Germany.

## Abstract

Respiratory viruses depend on host proteases for activation of viral fusion proteins, making these enzymes attractive targets for broad-spectrum antiviral strategies. We previously identified Trypstatin, a human Bikunin-derived Kunitz domain, as a potent endogenous inhibitor of the airway serine protease TMPRSS2. Here, we investigated whether TMPRSS2 inhibition is shared by additional human Kunitz domains. Kunitz domains with high sequence similarity to Trypstatin were synthesized, refolded, and functionally characterized. Domains derived from amyloid precursor protein (APP) and amyloid precursor-like protein 2 (APLP2) potently inhibited TMPRSS2, with APP displaying subnanomolar activity comparable to camostat mesylate. APP and APLP2 selectively blocked SARS-CoV-2 Spike-mediated entry without affecting VSV-G-mediated entry or cell viability and inhibited infection by multiple coronaviruses and influenza viruses, but not TMPRSS2-independent rhinovirus. In primary human airway epithelial cultures, APP and Trypstatin reduced replication of SARS-CoV-2, endemic coronaviruses, and influenza A virus, and remained stable in airway mucus. These findings identify APP and APLP2 Kunitz domains as potent endogenous inhibitors of TMPRSS2-dependent respiratory virus infection and promising scaffolds for host-directed broad-spectrum antivirals.

## Introduction

Respiratory viruses remain a major global health challenge, causing substantial morbidity, mortality, and economic burden worldwide. Seasonal pathogens such as influenza viruses, coronaviruses, and respiratory syncytial virus (RSV) are responsible for recurrent outbreaks, while the emergence of novel zoonotic viruses, exemplified by severe acute respiratory syndrome coronavirus 2 (SARS-CoV-2), highlights the continuing pandemic potential of respiratory infections. Although direct-acting antivirals have improved treatment options for selected viruses, their efficacy is frequently compromised by the rapid evolution of viral genomes and the emergence of drug-resistant variants. Consequently, there is increasing interest in host-directed antiviral strategies that target cellular factors required for viral replication and are therefore less susceptible to viral escape.

A common feature of many enveloped respiratory viruses is their dependence on host proteases for activation of viral fusion proteins. Among these, the type II transmembrane serine protease TMPRSS2 has emerged as a key host factor for viral entry^1–3^. TMPRSS2 cleaves and activates the Spike proteins of coronaviruses, including SARS-CoV-1, SARS-CoV-2, and endemic human coronaviruses^4,5^, as well as the hemagglutinin proteins of influenza A and B viruses^3^. Genetic and experimental studies have demonstrated that TMPRSS2 is critical for efficient viral replication and pathogenesis *in vivo*^6,7^. Because multiple genetically distinct respiratory viruses converge on TMPRSS2-dependent activation, this protease represents an attractive target for broad-spectrum antiviral intervention.

The therapeutic potential of targeting host proteases has been demonstrated by several naturally occurring protease inhibitors^1,8^. The bovine Kunitz inhibitor aprotinin inhibits host proteases involved in viral glycoprotein activation and exhibits antiviral activity against influenza viruses and coronaviruses^9,10^. Likewise, the endogenous serpin α1-antitrypsin inhibits TMPRSS2 and suppresses SARS-CoV-2 infection, while antithrombin has recently been reported to inhibit TMPRSS2-dependent viral infection through related mechanisms^11^. Together, these findings suggest that endogenous protease inhibitors constitute an important but incompletely explored component of antiviral host defense and may provide attractive scaffolds for the development of host-directed antiviral therapeutics.

Among endogenous protease inhibitors, Kunitz-type inhibitors represent a structurally conserved family characterized by a compact fold stabilized by three disulfide bridges and a reactive protease-binding loop that mediates high-affinity interactions with target enzymes^12,13^. Kunitz domains occur either as standalone proteins or as modules embedded within larger multidomain precursors and regulate diverse physiological processes^14^. Several human proteins contain Kunitz domains, including bikunin^15^, amyloid precursor protein (APP)^16^, amyloid precursor-like proteins (APLPs)^17^, WAP, follistatin/Kazal, immunoglobulin, Kunitz, and netrin-domain containing protein (WFIKKN) protein^18^, and collagen α3(VI)^19^. While these proteins have been implicated in coagulation, inflammation, extracellular proteolysis, and tissue homeostasis, their potential contribution to antiviral host defense remains largely unexplored^20^.

Recently, we identified Trypstatin, a C-terminal Kunitz domain derived from the human protease inhibitor bikunin, as a potent endogenous inhibitor of TMPRSS2 and related airway proteases^21^. Trypstatin efficiently inhibited infection by coronaviruses and influenza viruses in vitro and reduced viral replication *in vivo*, demonstrating that endogenous human Kunitz domains can function as host-directed antiviral agents. These findings raised the question of whether antiviral TMPRSS2 inhibition is a unique property of Trypstatin or a more broadly shared feature among human Kunitz domains.

To address this question, we systematically investigated human Kunitz domains with high sequence similarity to Trypstatin for their ability to inhibit TMPRSS2 and block respiratory virus infection. We identified several endogenous TMPRSS2 inhibitors, including Kunitz domains derived from APP and APLP2, that efficiently inhibited TMPRSS2 activity and suppressed infection by multiple coronaviruses and influenza viruses. Our findings demonstrate that antiviral TMPRSS2 inhibition is not unique to Trypstatin but is shared by multiple human Kunitz domains, identifying APP and APLP2 as potent endogenous inhibitors of respiratory virus infection and revealing Kunitz-domain-containing proteins as a previously underappreciated source of host-directed antiviral molecules.

## Methods

### Cell culture

Caco-2 cells (human colorectal adenocarcinoma cells, kindly provided by Prof. Holger Barth, Ulm University), HEK293T cells (ATCC #CRL-3216), Calu-3 cells (ATCC #HTB-55), LLC-MK2 cells (kindly provided by Lia van der Hoek, University of Amsterdam), and Vero E6 cells (ATCC #CRL-1586) were maintained at 37 °C in a humidified atmosphere containing 5% CO₂. Caco-2, HEK293T, H1Hela and Vero E6 cells were cultured in Dulbecco’s Modified Eagle Medium (DMEM), whereas Calu-3 and LLC-MK2 cells were maintained in Minimum Essential Medium Eagle (MEM). Media were supplemented with fetal calf serum (FCS), penicillin (100 U/mL), streptomycin (100 μg/mL), 2 mM L-glutamine, 1 mM sodium pyruvate, and 1x non-essential amino acids as required for the respective cell line. Cells were routinely tested and confirmed negative for mycoplasma contamination.

### Generation of human airway epithelial cells

Differentiated air-liquid interface cultures of human airway epithelial cells (HAECs) were generated from primary human basal cells isolated from airway epithelia as described previously^11^. In brief, cells were expanded in Airway Epithelial Cell Basal Medium supplemented with Airway Epithelial Cell Growth medium SupplementPack (PromoCell). Medium was replaced every 2 days until 90 % confluency was reached and the HAECs were detached using DetachKIT (PromoCell) and seeded into 6.5 mm Transwell filters (Corning Costar). Filters were pre-coated with Collagen Solution (StemCell Technologies) overnight and irradiated with UV light for 30 min before 35,000 cells were seeded onto the apical side of each filter in 200 µl of growth medium supported by 600 µl of growth medium in the basolateral part. After 72-96 hours, when the cells reached confluence, apical medium was removed and basal medium was replaced by differentiation medium consisting of a 1:1 mixture of DMEM-H and LHC Basal (Thermo Fisher) supplemented with Airway Epithelial Growth Medium SupplementPack. Medium was replaced every 2 days. Removal of apical medium (air lifting) defined day 0 of air-liquid interface (ALI) culture. Cells were grown under ALI conditions until experiments were performed at day 25-27.

### Peptide synthesis and refolding

Kunitz domain sequences corresponding to Trypstatin (AMBP 284-344, UniProt P02760), WFIKKN1 domain 3 (Q96NZ8), WFIKKN2 domain 3, amyloid precursor protein (APP, P05067), amyloid precursor-like protein 2 (APLP2, Q06481), and collagen α-3(VI) chain (P12111) were retrieved from UniProt. For WFIKKN1, WFIKKN2, APP, APLP2, and collagen α-3(VI), an additional glycine residue was introduced at both the N- and C-terminus to facilitate peptide synthesis. Peptides were synthesized by Synpeptide Co. Ltd. (Shanghai, China). To enable disulfide bond formation, peptides were subjected to oxidative refolding. Briefly, 10 mg of peptide was dissolved in 4 mL denaturing buffer supplemented with DTT (1 mM) and incubated for 40 min at 40 °C. Subsequently, 4 mL of denaturing buffer (50 mM Tris pH 8.5, 157 mM NaCl, 6 M GuHCl) and glutathione disulfide (GSSG; 59.8 mg) were added, and the mixture was dialyzed against 500 mL renaturing buffer (50 mM Tris pH 8.5, 157 mM NaCl) for 16 h at room temperature under continuous stirring.

Refolded peptides were purified by reversed-phase high-performance liquid chromatography (RP-HPLC) using a C18 column (10 × 250 mm, 5 μm particle size) at a flow rate of 3 mL/min. Separation was performed using a linear gradient (min/%B): 0/5, 5/15, 65/45, and 75/80, with UV detection at 280 nm. B being 0.1 % TFA in acetonitrile. Fractions were collected at one-minute intervals and dried using a vacuum concentrator. Anti-protease activity was detected exclusively in the dominant chromatographic peak, whereas minor peaks showed no detectable activity.

Correct folding and disulfide bond formation were verified by matrix-assisted laser desorption/ionization time-of-flight (MALDI-TOF) mass spectrometry using an Axima Confidence instrument (Shimadzu) operated in positive linear mode. Samples were spotted onto a 384-well stainless-steel target plate pre-coated with α-cyano-4-hydroxycinnamic acid (CHCA; 5 mg/mL). Equal volumes (0.5 μL) of sample and matrix solution were mixed directly on the target and air-dried prior to analysis. Spectra were acquired using a 337 nm nitrogen laser at an acceleration voltage of 20 kV, with 100 profiles collected per sample (20 laser shots per profile). External calibration was performed using the TOFMix™ MALDI calibration kit. Data acquisition and analysis were conducted using Shimadzu Biotech Launchpad software (version 2.9.8.1).

Fractions exhibiting the expected molecular mass corresponding to correctly folded Kunitz domains (consistent with three disulfide bridges) were selected for downstream assays.

### RP-HPLC analysis of peptides in mucus

A 15 µL aliquot of each sample was diluted with 75 µL of 5% acetonitrile/0.1% TFA in water and subjected to reversed-phase chromatographic analysis on a Biobasic C18 RP-HPLC column (Thermo Scientific, Waltham, MA, USA) with dimensions of 2.1 × 100 mm (5 µm). The separation was performed at 0.5 mL/min using the gradient 0/5, 10/45, and 12/100 (total running time in min/%B), with A being 0.1% TFA in water and B being 0.1% TFA in acetonitrile. All separations were run on an Agilent 1100 HPLC system (Agilent Technologies, Santa Clara, CA, USA). Online UV detection was performed at 225 nm. Chromatogram recording and peak integration were done with ChemStation B.04.03 (Agilent Technologies, Santa Clara, CA, USA).

### MALDI-TOF analysis of peptides in mucus

The samples were analyzed with an Axima Confidence MALDI-TOF MS (Shimadzu, Kyoto, Japan) in positive reflector mode on a 384-spot stainless-steel sample plate. Spots were coated with 0.5 µL of 10 mg/mL CHCA dissolved in a mixture of TFA/water/2-propanol/acetonitrile (2.5/47.5/25/25), and the solvent was allowed to air-dry. Samples were mixed with 4 volumes of 2.5% TFA, and 0.5 µL of each diluted sample was applied to the dry, pre-coated well and immediately mixed with 0.5 µL of matrix; the mixture was allowed to air-dry. Ten replicates were applied per sample. All spectra were acquired in the positive ion linear mode using a 337-nm N2 laser. An accelerating voltage of 20 kV was applied to the ion source. Laser shots were automatically performed using a circular raster with a diameter of 2000 μm and a spacing of 200 μm for each well; 100 profiles were acquired per sample, with 20 shots accumulated per profile. The equipment was calibrated with a standard mixture provided in the TOFMixTM MALDI kit (Shimadzu, Kyoto, Japan). Measurements and MS data processing were controlled by the MALDI-MS Application Shimadzu Biotech Launchpad 2.9.8.1 (Shimadzu, Kyoto, Japan). Data processing was performed with GraphPad Prism version 10.3.1 for Windows (GraphPad Software, Boston, MA, USA).

### Generation of lentiviral pseudoparticles

Lentiviral pseudoparticles bearing coronavirus spike proteins were generated by seeding 9 × 10⁵ HEK293T cells in 2 mL of culture medium in 6-well plates. On the following day, cells were co-transfected with 0.49 µg pCMVdR8_91 (encoding a replication-deficient lentiviral backbone), 0.49 µg pSEW-Luc2 (encoding a luciferase reporter gene; both kindly provided by Christian Buchholz, Paul-Ehrlich-Institute), and 0.02 µg of the respective glycoprotein expression plasmid. The latter included pCG1_SARS-2-SΔ18 (SARS-CoV-2 Omicron XBB.1.5; kindly provided by Stefan Pöhlmann, German Primate Center), pCG1_MERS-CoV, or pCG1_SARS-CoV-1 (both kindly provided by Michael Schindler, University of Tübingen). Plasmid DNA was combined with TransIT®-LT1 transfection reagent at a 1:3 ratio in serum-free medium and incubated for 20 min at room temperature prior to addition. The transfection mixture was then added dropwise to the cells. At 48 h post-transfection, supernatants containing pseudoparticles were collected and clarified by centrifugation at 1,500 rpm for 5 min. Virus-containing supernatants were subsequently aliquoted and stored at −80 °C until further use.

### Pseudoparticle inhibition assay

For transduction experiments, 1 × 10⁴ Caco-2 cells were seeded in a 96-well flat-bottom plate one day prior to infection. Cells were maintained in Dulbecco’s Modified Eagle Medium (DMEM) supplemented with 10% fetal calf serum, 2 mM L-glutamine, 100 U/mL penicillin, 100 µg/mL streptomycin, 1× non-essential amino acids, and 1 mM sodium pyruvate. On the day of transduction, the culture medium was replaced with 60 µL of serum-free medium. Cells were pre-treated with serial dilutions of test compounds for 30 min at 37 °C, followed by the addition of 20 µL of the respective lentiviral pseudoparticles. Transduction efficiency was determined 48 h post-transduction by measuring luciferase activity in cell lysates using the Luciferase Assay System (Promega). Luminescence was recorded with an Orion II microplate reader using Simplicity 4.2 software. Values obtained for H₂O-treated controls were defined as 100% pseudoparticle entry.

### Viral strains and propagation

Human coronavirus NL63 (hCoV-NL63) was obtained from Lia van der Hoek, Amsterdam University, hCoV-229E was obtained from ATCC (#VR-740) and propagated as described before^22,23^. SARS-CoV-2 Omicron BA.5 (B.1.1.529) was kindly provided by Prof. Dr. Florian Schmidt (University of Bonn) and propagated in Caco-2 cells. hRV16 was obtained from ATCC (#VR-283). In brief, H1Hela cells were inoculated with an MOI of 0.1. Two days post-infection, virus-containing supernatant was harvested and cleared by centrifugation. Influenza A/Virginia/ATCC3/2009 (Virg09; A/H1N1 subtype) was purchased from ATCC (VR-1737), influenza A/Victoria/361/2011 virus (Vict11; A/H3N2) was donated by G. Rimmelzwaan (Rotterdam, The Netherlands), influenza B/Malaysia/2506/2004 (B/Mal; B/Victoria-lineage) was obtained through the NIH Biodefense and Emerging Infections Research Resources Repository, NIAID, NIH (BEI Resources, NR-12280).

### Infection of human airway epithelial cells (HAECs)

Right before infection, the apical surface of HAECs grown on Transwell filters at the air-liquid interface were washed three times with pre-warmed PBS to remove mucus. Subsequently, 10 µM Trypstatin, 10 µM camostat mesylate, 10 µM APP or PBS were added onto the apical surface for 10 minutes before viral inoculum (SARS-CoV-2 Omicron BA.5 MOI 0.5, hCoV-229E MOI 0.05, hCoV-NL63 MOI 0.05, IAV H1N1 PR8 MOI 0.05) was added for two hours. Following this, the inoculum and compound was removed, cells washed with pre-warmed PBS and further cultured at the air-liquid interface. Accumulated mucus was washed off daily with pre-warmed PBS. Two (three days for hCoV-NL63) days post-infection, mucus was removed and PBS added to the apical side for 30 minutes before the sample was subjected to TCID_50_ titration on the respective permissive cell line. TCID_50_ was calculated 6 days post-infection according to Reed and Muench^24^.

### hCoV-NL63 inhibition assay

2.5 × 10⁴ Caco-2 cells were seeded in a 96-well flat-bottom plate one day prior to infection. Cells were pre-treated with serial dilutions of test compounds in phosphate-buffered saline (PBS) for 30 min, followed by infection with hCoV-NL63 at a multiplicity of infection (MOI) of 0.018. Infected cells were incubated at 33 °C. At 48 h post-infection (hpi), cells were detached and fixed with 4% paraformaldehyde (PFA) for 30 min. After fixation, cells were washed with PBS and stained for flow cytometric analysis using an anti-NL63 nucleocapsid antibody (Sino Biological, #40641-T62) diluted 1:5,000 in Buffer B (Fix & Perm, MuBio Nordic) for 40 min at 4 °C. Cells were subsequently washed twice with FACS buffer (1% fetal calf serum in PBS) and incubated with an Alexa Fluor 647-conjugated anti-rabbit secondary antibody (Cell Signaling Technology, #4414), diluted 1:5,000 in FACS buffer, for 30 min at 4°C. Following two additional washing steps with FACS buffer, samples were analyzed by flow cytometry using a CytoFLEX LX instrument (Beckman Coulter) and CytExpert 2.3 software.

### Human rhinovirus 16 inhibition assay

2 × 10^4^ H1HeLa-cells were seeded in a 96-well flat-bottom plate one day prior to infection. Cells were treated with serial dilution of compounds in PBS before inoculation with hRV16 at a MOI of 0.01. Two days post infection, cell viability was assessed by MTT assay. To this end, supernatant was removed and cells incubated in MTT solution for 2.5 h at 37°C. Afterwards, MTT was removed and a 1:1 mixture of EtOH/DMSO solution added onto the cells before measurement of absorption at 490 nm with a VersaMax microplate reader with SoftMax Pro 7 software.

### Influenza virus inhibition assay

Antiviral activity in Calu-3 cells infected with influenza A or B virus was assessed using an NP immunostaining-based high-content imaging assay, adapted from a previously described protocol^25^. Calu-3 cells were seeded in black 96-well plates at 30,000 cells per well and incubated overnight. The following day, cells were pretreated with serial dilutions of the compounds for 30 min before infection with influenza virus A/Virginia/ATCC3/2009 (Virg09), A/Victoria/361/2011 (Vict11), or B/Malaysia/2506/2004 (B/Mal), at 100 TCID50 per well. After three days at 35 °C, cells were fixed with 2% paraformaldehyde, permeabilized with 0.1% Triton X-100, and stained for viral NP (for IAV: ab20343 [Abcam] at 1:1000; for IBV: RIF17 R2/3 [HyTest] at 1:2000). Alexa Fluor 488 goat anti-mouse IgG (A11001 [Invitrogen] at 1:500) was used as the secondary antibody, and nuclei were counterstained with Hoechst (Thermo Fisher Scientific). Nine fields per well were acquired using a CellInsight CX5 high-content imaging platform (Thermo Fisher Scientific), and the percentage of virus-infected cells was calculated as the number of NP-positive cells relative to the total number of cells.

### Recombinant TMPRSS1, TMPRSS2 and TMPRSS14 activity assay

To assess inhibition of recombinant human transmembrane serine proteases (TMPRSS), serial dilutions of test compounds (25 µL) were incubated with 25 µL of recombinant enzyme in the appropriate assay buffer for 15 min at 37 °C. The following enzymes and concentrations were used: TMPRSS2 (2 µg/mL; LSBio, #LS-G57269), TMPRSS1 (0.02 µg/mL; R&D Systems, #11416), and TMPRSS14 (0.2 µg/mL; R&D Systems, #3946).

Assays for TMPRSS2 were performed in buffer containing 50 mM Tris-HCl and 0.154 mM NaCl (pH 8.0), whereas TMPRSS1 and TMPRSS14 assays were conducted in buffer containing 50 mM Tris and 0.05% Brij35 (pH 9.0). Following pre-incubation, 50 µL of fluorogenic protease substrate was added. For TMPRSS2 and TMPRSS14, 20 µM BOC-Gln-Ala-Arg-AMC (Bachem, #4017019) was used, whereas TMPRSS1 assays employed 150 µM BOC-Gln-Arg-Arg-AMC (PeptaNova). Reactions were incubated at 37 °C for 2 h (TMPRSS2) or 5 min (TMPRSS1 and TMPRSS14). Fluorescence intensity was measured at an excitation wavelength of 380 nm and an emission wavelength of 460 nm using a Synergy™ H1 microplate reader (BioTek) equipped with Gen5 3.04 software.

### Recombinant neutrophil elastase activity assay

Neutrophil elastase activity was determined by incubating 25 µL of test compound with 25 µL of recombinant Neutrophil elastase (2 ng/µL; Merck Millipore, #324681) in assay buffer (50 mM Tris, 1 M NaCl, 0.05% (w/v) Brij-35, pH 7.5) for 15 min at 37 °C. Subsequently, 50 µL of 200 µM fluorogenic substrate MEOSUC-Ala-Ala-Pro-Val-AMC (Bachem, #4005227) was added, and the reaction mixture was incubated at 37 °C. Fluorescence intensity was measured after 5 min at an excitation wavelength of 380 nm and an emission wavelength of 460 nm using a Synergy™ H1 microplate reader (BioTek) with Gen5 3.04 software.

### Recombinant kallikrein 5 activity assay

Kallikrein 5 activity was assessed by incubating 25 µL of test compound with 25 µL of recombinant Kallikrein 5 (2 µg/mL; R&D Systems, #1108) in assay buffer (100 mM NaH₂PO₄, pH 8.0) for 15 min at 37 °C. Following pre-incubation, 50 µL of the fluorogenic substrate BOC-Val-Pro-Arg-AMC (R&D Systems, #ES011) was added, and the reaction mixture was incubated for 5 min at 37 °C. Fluorescence intensity was then measured at an excitation wavelength of 380 nm and an emission wavelength of 460 nm using a Synergy™ H1 microplate reader (BioTek) equipped with Gen5 3.04 software.

### Cellular TMPRSS activity assay

For cell-based protease activity assays, 2 × 10⁴ HEK293T cells were seeded in a 96-well flat-bottom plate. On the following day, cells were transiently transfected with 100 ng per well of an expression plasmid (EF1α mammalian expression vector backbone; Twist Bioscience) encoding the respective transmembrane serine protease, including TMPRSS2 (human, NCBI RefSeq: NP_005647.3), TMPRSS11D (NCBI RefSeq: NP_004253.1), and TMPRSS13 (NCBI RefSeq: NP_001070731.1). Transfection was performed using TransIT®-LT1 reagent at a DNA-to-reagent ratio of 1:3 in serum-free medium. After 20 min incubation at room temperature, the transfection mixture was added dropwise to the cells. At 16 h post-transfection, the medium was replaced with serum-free medium, and cells were treated with serial dilutions of test compounds in PBS. Following a 15 min incubation at 37 °C, protease-specific fluorogenic substrates (100 µM final concentration) were added: Boc-QAR-AMC for TMPRSS2 (Bachem, #4017019), Boc-FSR-AMC for TMPRSS11D (Bachem, #4012340), and Pyr-RTKR-AMC for TMPRSS13 (Bachem, #4018149). Reactions were incubated for 2 h at 37 °C. Fluorescence intensity was measured at an excitation wavelength of 380 nm and an emission wavelength of 460 nm using a Synergy™ H1 microplate reader (BioTek) with Gen5 3.04 software. Measured values were background-corrected using non-transfected HEK293T cells.

### Cathepsin activity assay

Serial dilutions of test compounds were incubated with recombinant Cathepsin L (R&D Systems, #952-CY) in assay buffer (50 mM MES, 5 mM DTT, 1 mM EDTA, 0.005% Brij35, pH 6.0) or with Cathepsin B derived from human placenta (Sigma-Aldrich, #C0150) in assay buffer (105.6 mM KH₂PO₄, 14.4 mM Na₂HPO₄, 1.2 mM EDTA, 0.07% Brij35, 2.4 mM L-cysteine, pH 6.0). Protease–inhibitor mixtures were incubated for 10 min at room temperature prior to the addition of fluorogenic substrates. For cathepsin L, Z-Leu-Arg-AMC (Bachem, #4034611) was used, whereas cathepsin B assays employed Z-Arg-Arg-AMC (Bachem, #4004789). Final concentrations were 0.01 µg/mL for cathepsin L, 1 µg/mL for cathepsin B, and 40 µM or 30 µM for the respective substrates. Fluorescence intensity was recorded at an excitation wavelength of 380 nm and an emission wavelength of 460 nm using a Synergy™ H1 microplate reader (BioTek) with Gen5 3.04 software. Measurements were taken after 25 min incubation at 37 °C for cathepsin L or after 60 min at 40 °C for cathepsin B.

### Reagents

Camostat mesylate (#SML0057) and E-64d (#E8640) were obtained from Merck. Aprotinin (#A1153) and Rupintrivir (PZ0315-25MG) from Sigma Aldrich, and baloxavir acid from MedChemExpress (HY-109025A). Remdesivir (GS-5734) was obtained from Selleckchem.

### Cytotoxicity assay

To evaluate cytotoxic effects of test compounds, 1 × 10⁴ Caco-2 cells were seeded in a 96-well flat-bottom plate. On the following day, culture medium was replaced, and cells were treated with serial dilutions of compounds or PBS as a control. After 48 h of incubation, cell viability was determined by quantifying intracellular ATP levels using the CellTiter-Glo® assay (Promega), according to the manufacturer’s instructions. Luminescence was measured using an Orion II microplate reader with Simplicity 4.2 software.

### Statistical Analysis

Unless stated otherwise, analysis was performed using GraphPad Prism version 10.6.1. Calculation of IC_50_ values via nonlinear regression was performed using normalized response-variable slope equation. The number of repeated experiments is stated in each figure legend.

## Results

### Identification and synthesis of human Kunitz homologues to Trypstatin

Following the identification of Trypstatin as a potent endogenous inhibitor of TMPRSS2 with broad antiviral activity^21^, we asked whether antiviral TMPRSS2 inhibition is unique to Trypstatin or shared by additional human Kunitz domains. To address this question, we performed a systematic search of the MEROPS and UniProt databases, identifying 26 Kunitz domains encoded in the human genome (Figure 1A).

**Figure 1.**
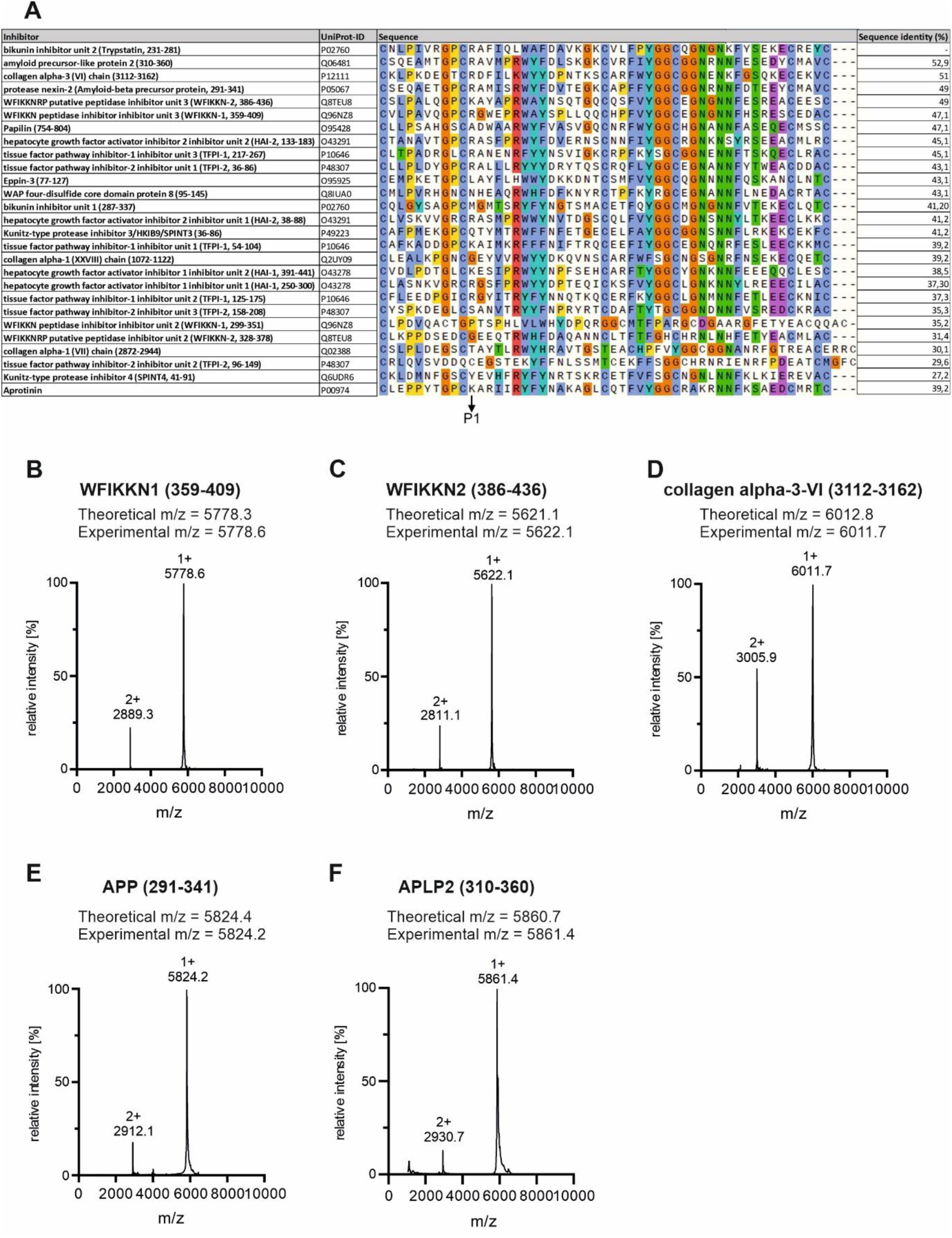
Identification, synthesis and structural validation of human Kunitz domains. (A) Human Kunitz domains were identified by searching the MEROPS and UniProt databases. Pairwise sequence alignments against Trypstatin were performed using the EMBOSS Needleman-Wunsch algorithm, and the five domains exhibiting the highest sequence identity were selected for experimental characterization. (B-F) Kunitz domains derived from APLP2 (aa 310-360), COL6A3 (aa 3112-3162), APP (aa 291-341), WFIKKN2 (aa 386-436), and WFIKKN1 (aa 359-409) were chemically synthesized and subjected to oxidative refolding. Refolded peptides were purified by reversed-phase HPLC and analyzed by MALDI-TOF mass spectrometry. Observed molecular masses were consistent with the expected masses of fully oxidized Kunitz domains containing three intramolecular disulfide bridges. Modified from Lawrenz, 2026^26^ (CC BY 4.0; https://creativecommons.org/licenses/by/4.0/).

Because sequence similarity is frequently associated with conservation of protease specificity among Kunitz inhibitors, pairwise sequence alignments against the Kunitz domain of Trypstatin were performed using the EMBOSS Needleman-Wunsch algorithm. The five domains exhibiting the highest sequence similarity were selected for experimental characterization. These included Kunitz domains derived from APLP2 (aa 310-360; 52.9 % identity), COL6A3 (aa 3112-3162; 51.0 % identity), APP (aa 291-341; 49.0 % identity), WFIKKN2 (aa 386-436; 49.0 % identity), and WFIKKN1 (aa 359-409; 47.1 % identity).

The selected domains were chemically synthesized and subjected to oxidative refolding to promote formation of the characteristic Kunitz-domain disulfide architecture. Refolded peptides were purified by reversed-phase HPLC and analyzed by MALDI-TOF mass spectrometry. For all candidates, a dominant chromatographic peak was obtained after refolding, whereas only minor amounts of alternative species were detected (Figure 1B-F).

Mass spectrometric analysis revealed molecular masses that closely matched the theoretical masses expected for the fully oxidized proteins, consistent with correct folding and formation of the expected three intramolecular disulfide bridges (Figure 1B-F). Collectively, these results established a panel of structurally validated human Kunitz domains for subsequent analyses of protease inhibition and antiviral activity.

### APP- and APLP2-derived Kunitz domains are potent and selective inhibitors of TMPRSS2

To determine whether the selected human Kunitz domains inhibit TMPRSS2, their anti-proteolytic activity was evaluated in fluorogenic assays using recombinant TMPRSS2. Trypstatin, camostat mesylate^4^, aprotinin^9^, and E64d^4^ served as reference inhibitors. Four of the five investigated Kunitz domains inhibited TMPRSS2 activity, whereas the Kunitz domain derived from collagen α-3-VI (COL6A3) was completely inactive (Figure 2A). Thus, high sequence similarity to Trypstatin alone was not sufficient to predict TMPRSS2 inhibition.

**Figure 2:**
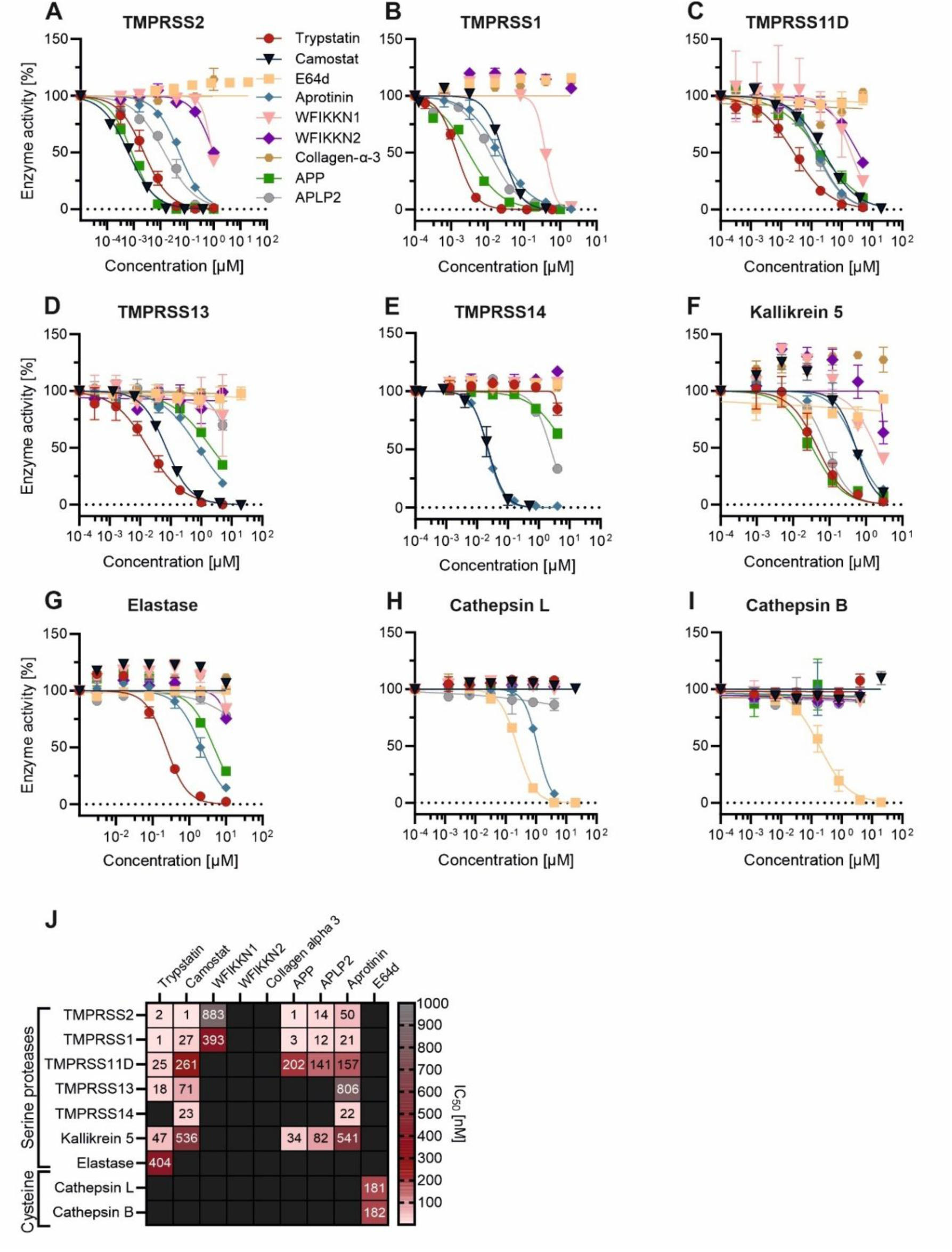
Anti-proteolytic profile of selected human Kunitz domains. **(A-B, F-I)** Serial dilutions of the indicated Kunitz domains and reference inhibitors pre-incubated with the indicated proteases prior to addition of a fluorogenic substrate. Camostat mesylate, aprotinin, and E64d served as reference inhibitors. Protease activity was quantified by measuring fluorescence (excitation: 380 nm; emission: 460 nm). **(C-D)** HEK293T cells were transfected with expression plasmids encoding TMPRSS11D, TMPRSS13, or an empty vector control. Cells were subsequently treated with the indicated inhibitors before addition of a fluorogenic substrate. Fluorescence signals were corrected for background activity derived from mock-transfected cells. Data represent mean values ± SEM from three independent experiments performed in technical triplicates. (J) Heatmap summarizing IC₅₀ values across all tested proteases. Black fields indicate IC₅₀ values >1000 nM. Reproduced from Lawrenz, 2026^26^ (CC BY 4.0; https://creativecommons.org/licenses/by/4.0/).

Among all candidates, the APP-derived Kunitz domain (KD) emerged as the most potent inhibitor. APP-KD inhibited TMPRSS2 with an IC_50_ value of 0.8 nM, exceeding the potency of Trypstatin (IC_50_ = 2.3 nM) and approaching that of the clinically used TMPRSS2 inhibitor camostat mesylate (0.6 nM). The closely related APLP2-derived KD also inhibited TMPRSS2 efficiently, albeit with lower potency (IC_50_ = 14.0 nM). In contrast, WFIKKN1- and WFIKKN2 KDs displayed weaker inhibition and did not suppress TMPRSS2 activity to baseline levels at the concentrations tested (Figure 2A). These findings identify APP KD as the most potent endogenous TMPRSS2 inhibitor examined in this study.

To further characterize their protease selectivity, the Kunitz domains were evaluated against a panel of proteases implicated in respiratory tract physiology and viral glycoprotein activation. Recombinant enzyme assays were performed for TMPRSS1, TMPRSS14, neutrophil elastase, and kallikrein 5, whereas TMPRSS11D and TMPRSS13 activity was assessed in cell-based assays using protease-transfected HEK293T cells (Figure 2B-G). APP KD displayed an inhibition profile that largely overlapped with that of Trypstatin and efficiently inhibited multiple members of the TMPRSS family as well as neutrophil elastase and kallikrein 5 (Figure 2B-G, J). However, quantitative differences between the inhibitors became apparent. While APP KD exhibited the strongest activity against TMPRSS2, Trypstatin generally showed greater potency against TMPRSS13, TMPRSS14, and neutrophil elastase (Figure 2J). APLP2 KD displayed a similar inhibition profile but was consistently less potent than APP KD and Trypstatin across most proteases examined.

Notably, WFIKKN1 and WFIKKN2 KDs retained substantial activity against TMPRSS2 and TMPRSS11D, despite exhibiting markedly reduced activity against TMPRSS1, TMPRSS13, TMPRSS14, neutrophil elastase, and kallikrein 5 (Figure 2B-G, J). Thus, although less potent than APP and APLP2 KDs, both proteins displayed a distinct and comparatively narrow protease inhibition spectrum. In contrast, COL6A3 KD remained inactive against all proteases tested (Figure 2J). Together, these findings demonstrate that closely related human KDs differ substantially in both potency and protease selectivity.

Because several respiratory viruses can utilize endosomal cysteine proteases as alternative entry factors^27,28^, we next assessed activity against cathepsin L and cathepsin B. None of the investigated human KDs exhibited detectable inhibition of either protease (Figure 2H,I). In contrast, the cysteine protease inhibitor E64d efficiently blocked both cathepsins, confirming assay performance^4,27^. These findings indicate that the inhibitory activity of the investigated KDs is largely restricted to serine proteases and does not extend to endosomal cysteine proteases involved in alternative viral entry pathways.

A comprehensive comparison of inhibitory activities is summarized in Figure 2J. Collectively, these results identify APP and APLP2 KDs as potent endogenous inhibitors of TMPRSS2 and related airway proteases. Notably, the APP-derived KD inhibited TMPRSS2 with potency comparable to camostat mesylate and approximately threefold greater than Trypstatin, highlighting APP as a previously unrecognized and highly potent endogenous inhibitor of this key respiratory virus host factor.

### APP and APLP2 KDs inhibit TMPRSS2-dependent SARS-CoV-2 Spike-mediated entry

To determine whether inhibition of TMPRSS2 translated into antiviral activity, the selected KDs were evaluated in a lentiviral pseudoparticle assay measuring SARS-CoV-2 Spike-mediated entry into Caco-2 cells. Vesicular stomatitis virus glycoprotein (VSV-G)-pseudotyped particles served as a control for TMPRSS2-independent viral entry, and cellular ATP levels were measured in parallel to assess potential cytotoxic effects.

Consistent with their anti-proteolytic activity against TMPRSS2, all KDs except COL6A3 inhibited SARS-CoV-2 Spike-mediated entry (Figure 3A). APP KD emerged as the most potent endogenous inhibitor, reducing viral entry with an IC_50_ value of 14.4 nM. This activity was comparable to that of camostat mesylate (IC_50_ = 17.4 nM) and exceeded the potency of Trypstatin (IC_50_ = 46.8 nM) and APLP2 KD (IC_50_ = 98.0 nM). WFIKKN1 and WFIKKN2 KDS also reduced Spike-mediated entry, albeit with lower efficacy, whereas COL6A3 KD was inactive in the entry assay.

**Figure 3:**
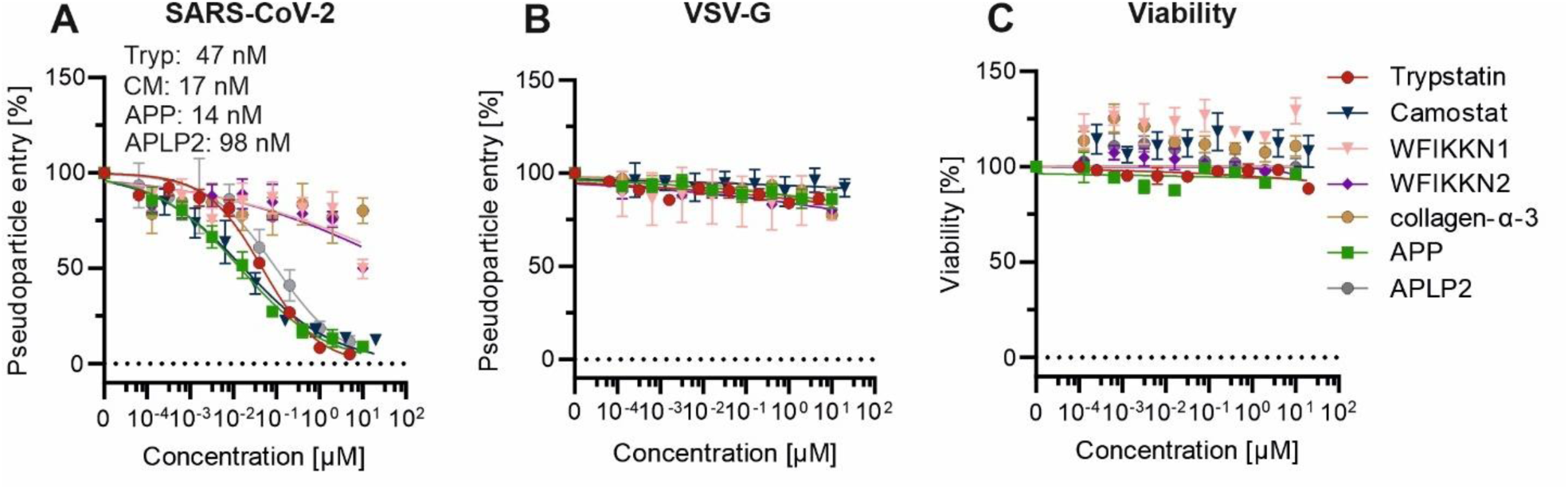
Selected human Kunitz domains inhibit SARS-CoV-2 Spike-mediated entry. **(A-B)** Caco-2 cells were pre-incubated with increasing concentrations of the indicated compounds prior to transduction with luciferase reporter lentiviral pseudoparticles pseudotyped with the SARS-CoV-2 Omicron XBB.1.5 Spike protein. Luciferase activity was quantified 48 h post-transduction as a measure of pseudovirus entry. (B) Specificity control using pseudoparticles pseudotyped with vesicular stomatitis virus glycoprotein (VSV-G), which enters cells independently of TMPRSS2. (C) Cytotoxicity analysis. Caco-2 cells were incubated with increasing concentrations of the indicated compounds for 48 h, followed by quantification of cellular ATP levels using the CellTiter-Glo® Luminescent Cell Viability Assay. Data represent mean ± SEM from three independent experiments performed in technical triplicates. Abbreviations: Tryp, Trypstatin; CM, camostat mesylate; APP, amyloid precursor protein; APLP2, amyloid precursor-like protein 2; WFIKKN, WAP, follistatin/Kazal, immunoglobulin, Kunitz, and netrin domain-containing protein. Reproduced from Lawrenz, 2026^26^ (CC BY 4.0; https://creativecommons.org/licenses/by/4.0/).

Importantly, the inhibitory potencies against SARS-CoV-2 Spike-mediated entry closely mirrored the corresponding TMPRSS2 inhibition profiles observed in Figure 2, supporting TMPRSS2 inhibition as the underlying mechanism of antiviral activity. Notably, APP KD was the most potent inhibitor in both assays, whereas COL6A3 KD was inactive in both protease and pseudovirus entry assay.

To assess specificity, the compounds were tested against VSV-G-mediated entry, which proceeds independently of TMPRSS2. None of the investigated KDs affected VSV-G pseudoparticle transduction, whereas SARS-CoV-2 Spike-mediated entry was efficiently inhibited (Figure 3B), arguing against nonspecific effects on lentiviral transduction or reporter gene expression.

Finally, none of the tested KDs reduced cellular ATP levels at concentrations exceeding those required to inhibit Spike-mediated entry (Figure 3C), indicating no detectable cytotoxicity. Together, these results demonstrate that APP and APLP2 KDs are potent and selective inhibitors of SARS-CoV-2 Spike-mediated entry and support TMPRSS2 inhibition and the underlying mechanism.

### APP and APLP2 KDs exhibit broad-spectrum antiviral activity against TMPRSS2-dependent respiratory viruses

Having established that selected human KDs inhibit TMPRSS2 and SARS-CoV-2 Spike-mediated entry, we next examined whether their antiviral activity extended to other TMPRSS2-dependent respiratory viruses. We first evaluated lentiviral pseudoparticles bearing the Spike proteins of SARS-CoV-1 or MERS-CoV, followed by experiments with authentic coronaviruses, influenza A and B viruses, and the TMPRSS2-independent rhinovirus HRV16.

APP KD, APLP2 KD, and Trypstatin inhibited SARS-CoV-1 Spike-mediated pseudoparticle entry into Caco-2 cells in a concentration-dependent manner (Figure 4A). APP KD showed the strongest inhibitory activity among the endogenous KDs (IC_50_ = 44 nM), consistent with its potency against TMPRSS2 and SARS-CoV-2 Spike-mediated entry (Figure 2A and 3A). APLP2 KD and Trypstatin were also active, demonstrating that these Kunitz domains inhibit entry mediated by Spike proteins from distinct sarbecoviruses.

**Figure 4:**
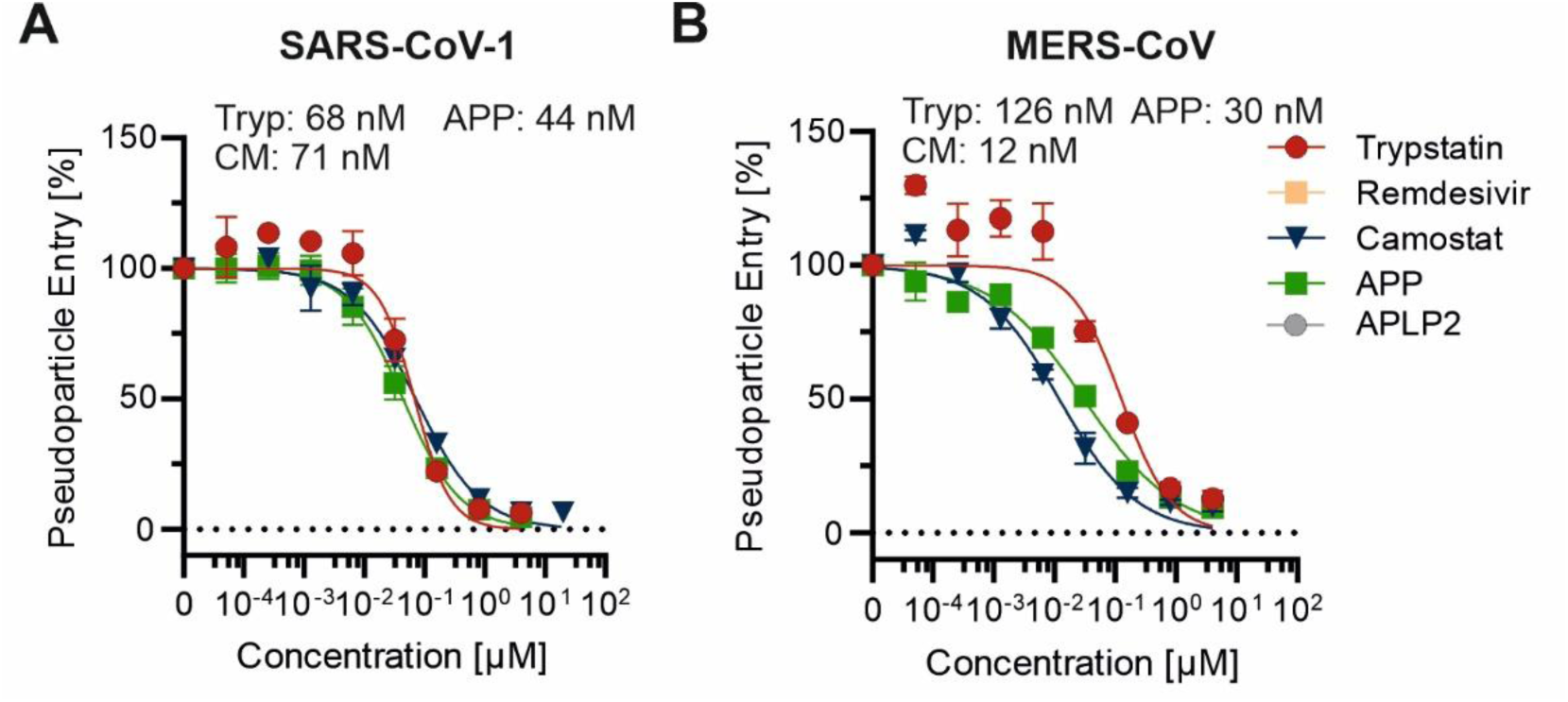
Human Kunitz domains inhibit SARS-CoV-1 and MERS-CoV Spike-mediated entry. Caco-2 cells were treated with serial dilutions of the indicated compounds before transduction with luciferase reporter lentiviral pseudoparticles pseudotyped with (A) SARS-CoV-1 Spike or (B) MERS-CoV Spike. Viral entry was quantified 48 h after transduction by measuring luciferase activity in cell lysates. Data represent mean ± SEM from three independent experiments performed in technical triplicates. Abbreviations: Tryp, Trypstatin; CM, camostat mesylate; APP, amyloid precursor protein; APLP2, amyloid precursor-like protein 2. Data were reproduced from Lawrenz *et al.*, 2026 under CC BY 4.0.

The compounds similarly inhibited entry mediated by MERS-CoV Spike (Figure 4B). APP KD again displayed pronounced activity (IC_50_ = 30 nM) compared to Trypstatin (IC_50_ = 126 nM). These results extend the antiviral activity of the Kunitz domains beyond SARS-related coronaviruses and demonstrate inhibition of Spike-mediated entry across all three highly pathogenic human coronaviruses tested: SARS-CoV-1, SARS-CoV-2, and MERS-CoV.

We next assessed the compounds against authentic hCoV-NL63 infection. In Caco-2 cells, APP KD, APLP2 KD, and Trypstatin inhibited hCoV-NL63 infection in a concentration-dependent manner, with IC_50_ values of 12, 108, and 17 nM, respectively (Figure 5A). APP KD was therefore the most potent inhibitor, followed closely by Trypstatin, whereas APLP2 KD showed lower but still substantial activity. The potency ranking closely matched that observed in the TMPRSS2 and pseudoparticle assays, supporting inhibition of TMPRSS2-dependent Spike activation as the principal mechanism of action.

**Figure 5:**
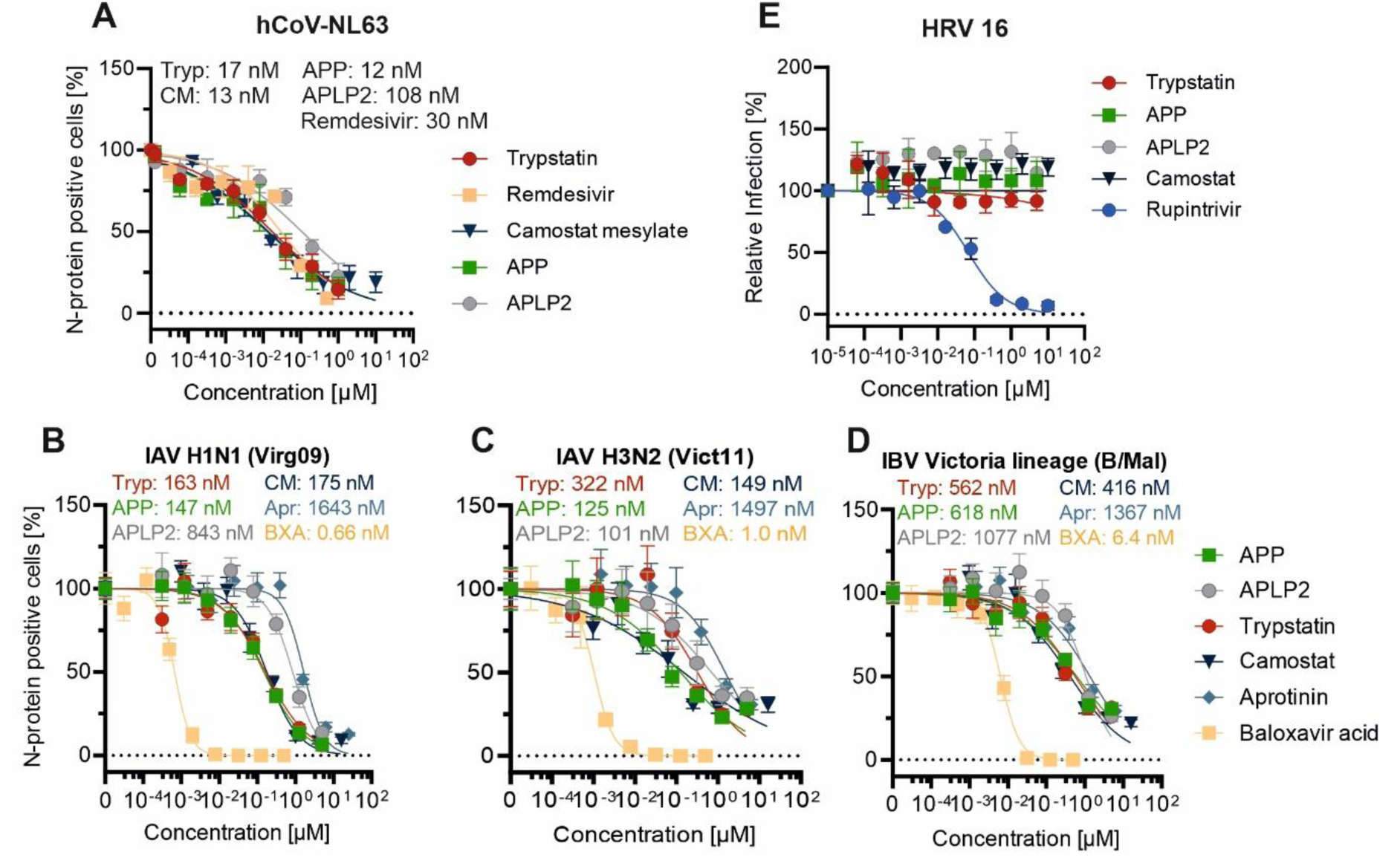
Human Kunitz domains inhibit authentic coronaviruses and influenza viruses but not HRV16. A) Caco-2 cells were incubated with increasing concentrations of the indicated compounds before infection with hCoV-NL63 at an MOI of 0.018. At 48 h post-infection, infected cells were quantified by flow-cytometric detection of viral nucleocapsid protein. (B-D) Calu-3 cells were exposed to serial dilutions of the indicated compounds for 30 min before inoculation with (B) influenza A virus H1N1 (A/Virginia/ATCC3/2009; Virg09), (C) influenza A virus H3N2 (A/Victoria/361/2011; Vict11), or (D) Victoria-lineage influenza B virus (B/Malaysia/2506/2004; B/Mal) at 100 TCID50 per well. Aprotinin and baloxavir acid (BXA) served as reference inhibitors. At three days post-infection, viral nucleoprotein-positive cells were quantified by immunostaining and high-content imaging. (E) H1HeLa cells were treated with serial dilutions of the indicated compounds before infection with HRV16 at an MOI of 0.01. Rupintrivir served as a positive-control inhibitor. At two days post-infection, cell viability was determined by MTT assay as a measure of protection against virus-induced cytopathic effects. Data represent mean ± SEM from three independent experiments. Abbreviations: Tryp, Trypstatin; APP, amyloid precursor protein; APLP2, amyloid precursor-like protein 2; BXA, baloxavir acid; hCoV-NL63, human coronavirus NL63; HRV16, human rhinovirus 16; IAV, influenza A virus; IBV, influenza B virus. Panel A was reproduced from Lawrenz et al., 2026 under CC BY 4.0.

We next investigated whether the antiviral activity of APP and APLP2 KDs extended to influenza viruses, which depend on host serine proteases for proteolytic activation of hemagglutinin. Aprotinin, a Kunitz-type inhibitor of extracellular serine proteases, and baloxavir acid (BXA), a direct-acting inhibitor of the influenza virus polymerase acidic protein endonuclease, served as reference inhibitors. APP KD, APLP2 KD, and Trypstatin inhibited influenza A virus H1N1 infection in Calu-3 cells in a concentration-dependent manner (Figure 5B). Aprotinin and BXA also efficiently inhibited infection, confirming the responsiveness of the assay to host protease- and virus-directed antiviral agents.

The Kunitz domains were similarly active against influenza A virus H3N2 (Figure 5C) and a Victoria-lineage influenza B virus (Figure 5D). Aprotinin and BXA also inhibited both viruses and served as positive controls. Across the three influenza viruses, APP KD and Trypstatin consistently showed high potency, with IC_50_ values ranging from 125 to 618 nM and from 163 to 562 nM, respectively. APLP2 KD was less potent, with IC_50_ values ranging from 701 to 1,077 nM. Together, these results demonstrate that the antiviral activity of APP- and APLP2-derived Kunitz domains extends to influenza A and B viruses. The activity of aprotinin further supports inhibition of host serine protease-mediated hemagglutinin activation as an effective antiviral mechanism, whereas the activity of BXA validates inhibition through an independent, virus-directed mechanism.

Finally, we examined whether the Kunitz domains affected HRV16, a non-enveloped rhinovirus that does not depend on TMPRSS2-mediated glycoprotein activation. Rupintrivir, a direct-acting inhibitor of the rhinovirus 3C protease, served as a positive control and protected cells against HRV16-induced cytopathic effects (Figure 5E). In contrast, none of the investigated Kunitz domains showed significant antiviral activity against HRV16. The activity of rupintrivir confirmed assay performance, while the inactivity of the Kunitz domains argues against nonspecific antiviral or cytoprotective effects and further supports inhibition of host serine protease-dependent viral activation as their principal mechanism of action.

Collectively, these findings demonstrate broad-spectrum activity of APP KD, APLP2 KD, and Trypstatin against diverse TMPRSS2-dependent respiratory viruses. Their activity against coronavirus Spike-mediated entry and authentic coronavirus and influenza virus infection, together with their lack of activity against HRV16, supports targeted inhibition of host protease-dependent viral activation.

### APP KD and Trypstatin inhibit respiratory virus replication in primary human airway epithelial cultures

To assess antiviral activity in a physiologically relevant airway model, we next evaluated the two most potent Kunitz domains, APP and Trypstatin, in differentiated primary human airway epithelial cells cultured at the air–liquid interface. The apical surface of human airway epithelial cultures from three independent donors was treated with APP, Trypstatin, camostat mesylate, or PBS prior to infection with SARS-CoV-2 Omicron BA.5, hCoV-229E, hCoV-NL63, or influenza A virus H1N1. Viral replication was quantified by titration of apical washes collected two or three days post-infection, depending on the virus.

Both APP KD and Trypstatin reduced replication of SARS-CoV-2 Omicron BA.5 in primary airway cultures from all donors tested, with antiviral effects comparable to or exceeding those observed for camostat mesylate (Figure 6A). Inhibition was also observed against the endemic coronaviruses hCoV-229E and hCoV-NL63, although the magnitude of reduction varied between donors and viruses (Figure 6B-C).

**Figure 6:**
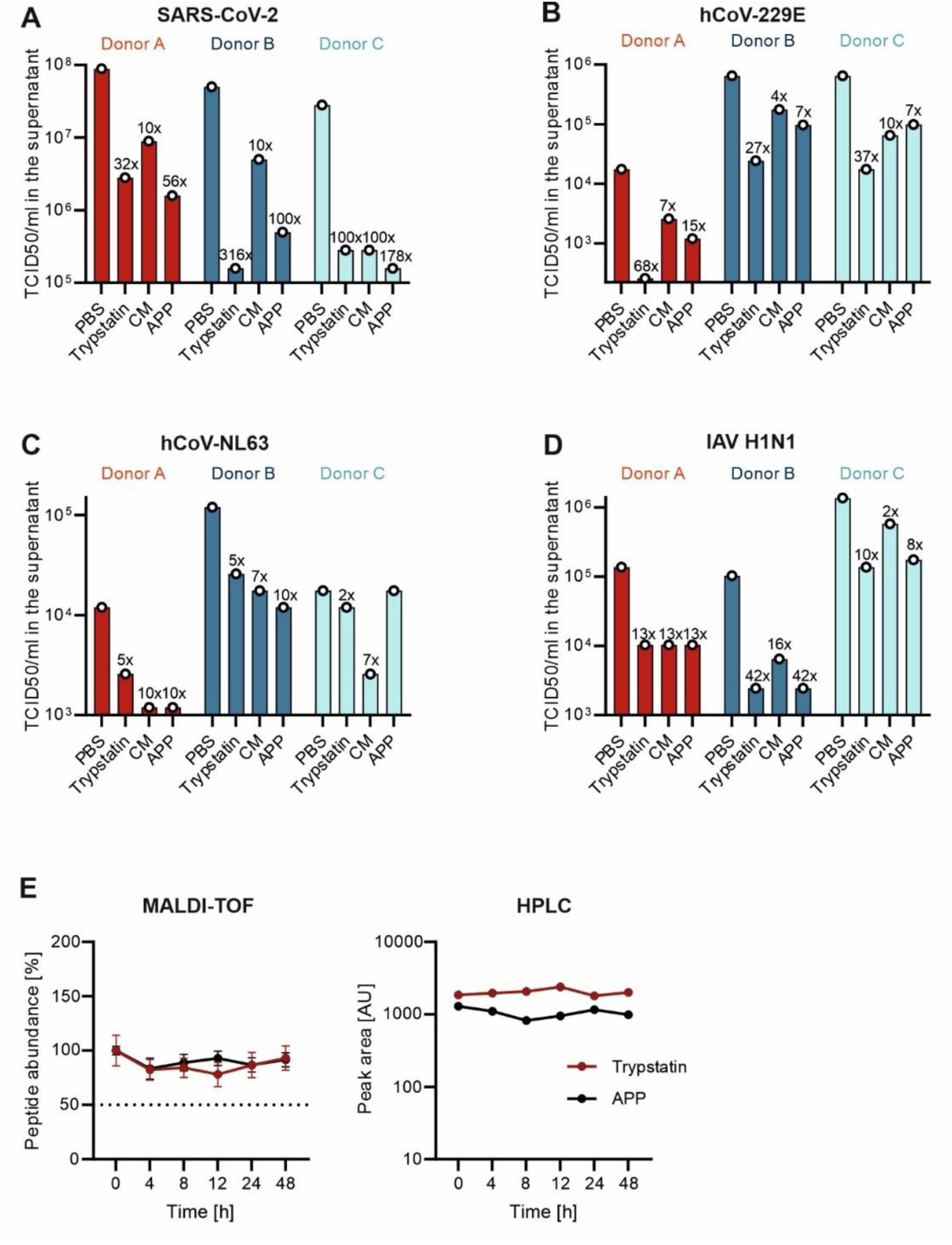
APP KD and Trypstatin inhibit different respiratory viruses in primary airway epithelium at the air-liquid-interface and remain stable in mucus over extended periods of time. **(A-D)** The apical site of human airway epithelial cells (HAEC) grown at the air–liquid interface was exposed to PBS, Trypstatin (10 µM), APP_KD (10 µM) or camostat mesylate (10 µM) before inoculation with SARS-CoV-2 Omicron BA. 5 (MOI 0.5), hCoV-NL63 (MOI 0.05), hCoV-229E (MOI 0.05) or IAV H1N1 PR8 (MOI 0.05) for 2 h before apical washing and further culturing at the air–liquid interface. Two (three for NL63) days-post infection, mucus was washed off and PBS was added to the apical side for 30 min before the sample was subjected to TCID_50_ titration. Shown are TCID_50_/ml values of pooled apical washes from two cultures per donor. Numbers above the bars indicate the fold reduction in infectious virus titer relative to the corresponding PBS-treated control for each donor **(E)** Respective peptide was incubated in mucus from HAECs at 37°C for the indicated time before analysis by MALDI-TOF (left) and HPLC (right).

APP KD and Trypstatin also reduced infectious influenza A H1N1 virus release in airway epithelial cultures, with donor-dependent differences in efficacy (Figure 6D). Together with the coronavirus data, these results indicate that APP KD and Trypstatin inhibit replication of distinct TMPRSS2-dependent respiratory viruses in primary airway tissue. These findings extend the activity observed in cell lines to a differentiated human airway epithelium model and support the relevance of Kunitz-mediated protease inhibition at the respiratory mucosal surface.

Since antiviral activity at the airway surface requires stability in mucus, we further examined the persistence of APP KD and Trypstatin in mucus derived from human airway epithelial cultures. Both peptides remained detectable over 48 h of incubation at 37 °C, as determined by MALDI-TOF mass spectrometry and RP-HPLC analysis (Figure 6E). Thus, APP KD and Trypstatin retain stability in a physiologically relevant mucus environment over extended periods of time.

Collectively, these findings demonstrate that APP KD and Trypstatin inhibit replication of coronaviruses and influenza virus in primary human airway epithelial cultures and remain stable in airway mucus, supporting their potential as endogenous host-directed antiviral inhibitors at the respiratory mucosal surface.

## Discussion

Host-directed antiviral strategies offer an attractive approach to overcome the limitations of virus-specific therapeutics, particularly for respiratory viruses that rapidly evolve and repeatedly cross species barriers. In this study, we identify human Kunitz domains as potent endogenous inhibitors of TMPRSS2-dependent respiratory virus infection. Building on the previous identification of Trypstatin as an endogenous TMPRSS2 inhibitor^21^, we show that antiviral protease inhibition is not restricted to this Bikunin-derived fragment but is shared by additional human Kunitz domains (KDs). In particular, KDs derived from APP and APLP2 potently inhibited TMPRSS2, blocked coronavirus Spike-mediated entry, and suppressed infection by multiple respiratory viruses. These findings reveal APP and APLP2 KDs as previously underappreciated endogenous inhibitors of respiratory virus infection and expand the concept of Kunitz domains as a source of host-directed antiviral molecules.

Among the KDs analyzed, APP KD emerged as the most potent TMPRSS2 inhibitor. Its inhibitory activity was comparable to camostat mesylate and exceeded that of Trypstatin in recombinant TMPRSS2 assays. This high potency translated into efficient inhibition of SARS-CoV-2 Spike-mediated entry, where APP KD again displayed the strongest activity among the endogenous domains tested. APLP2 KD also inhibited TMPRSS2 and viral entry, although with lower potency than APP KD. The close correlation between enzymatic TMPRSS2 inhibition and antiviral efficacy supports the conclusion that blockade of host protease activity is the primary mechanism underlying the antiviral effects of these peptides.

Despite their shared Kunitz scaffold and sequence similarity to Trypstatin, the selected domains differed substantially in potency and protease selectivity. APP and APLP2 KDs showed broad activity against TMPRSS2 and related airway serine proteases, whereas WFIKKN1 and WFIKKN2 KDs displayed weaker and more restricted inhibition profiles. In contrast, the KD of COL6A3 was inactive against all proteases tested, despite its high sequence similarity to Trypstatin. These findings indicate that sequence identity alone is insufficient to predict antiviral protease inhibition and suggest that specific residues within the reactive loop and surrounding interaction surface determine TMPRSS2 recognition. Structural studies of APP or APLP2 KDs bound to TMPRSS2 will be important to define the molecular determinants of this interaction and may guide rational engineering of optimized Kunitz-based inhibitors.

The inhibitory activity of APP and APLP2 KDs was largely restricted to serine proteases and did not extend to the cysteine proteases cathepsin L and B. This selectivity is relevant because coronaviruses can use either cell-surface serine proteases or endosomal cathepsins for entry, depending on the virus, cell type, and experimental system^28^. The lack of cathepsin inhibition, together with the absence of activity against VSV-G-mediated entry and TMPRSS2-independent rhinovirus infection, supports a specific mechanism targeting serine protease-dependent viral activation rather than nonspecific interference with viral entry, cellular viability, or reporter readout. Thus, APP and APLP2 KDs appear to act at a defined host-cell entry step shared by several respiratory viruses.

The antiviral activity of APP and APLP2 KDs extended beyond SARS-CoV-2. Both domains inhibited entry mediated by Spike proteins from SARS-CoV-1 and MERS-CoV, and APP, APLP2, and Trypstatin blocked authentic HCoV-NL63 infection. In addition, APP KD and Trypstatin inhibited influenza virus infection, consistent with the dependence of influenza hemagglutinin activation on airway serine proteases^29^. These data support the broader relevance of Kunitz-mediated protease inhibition against phylogenetically distinct respiratory viruses that converge on host protease-dependent glycoprotein activation. In contrast, the absence of inhibition against rhinovirus further strengthens the conclusion that antiviral activity is linked to TMPRSS2- or related serine protease-dependent entry mechanisms.

A key finding of this study is that APP KD and Trypstatin retained antiviral activity in differentiated primary human airway epithelial cultures grown at the air–liquid interface. This model more closely reflects the architecture and mucus-covered surface of the human respiratory epithelium than transformed cell lines^30^. APP KD and Trypstatin reduced replication of SARS-CoV-2 Omicron BA.5, endemic coronaviruses, and influenza A virus in cultures from independent donors, although the magnitude of inhibition varied between viruses and donors. Such donor-dependent variation is expected in primary airway models and may reflect differences in protease expression, mucus composition, epithelial differentiation, or innate antiviral responses. Importantly, the activity observed in this physiologically relevant system supports the potential relevance of Kunitz-domain-mediated protease inhibition at the respiratory mucosal surface.

The stability of APP and Trypstatin in airway mucus further supports their potential as antiviral scaffolds. Peptide and protein therapeutics intended for topical airway application must remain stable in a protease-rich mucosal environment. Both APP and Trypstatin remained detectable after prolonged incubation in mucus at 37 °C, as shown by MALDI-TOF and HPLC analyses. This suggests that the compact disulfide-stabilized Kunitz fold confers sufficient stability under conditions relevant to the respiratory tract. Together with their nanomolar potency, this property makes Kunitz domains attractive candidates for further development as locally applied host-directed antivirals.

The identification of APP and APLP2 KDs as potent TMPRSS2 inhibitors broadens the potential physiological functions of the APP protein family beyond its established roles in the nervous system. APP and APLP2 are type I transmembrane glycoproteins with large extracellular regions, short cytoplasmic tails, and multiple alternatively spliced isoforms^31–33^. In APP, the 751- and 770-amino-acid isoforms contain the Kunitz protease inhibitor (KPI) domain, whereas the predominantly neuronal APP695 isoform lacks it. APLP2 likewise occurs as KPI-containing and KPI-deficient splice variants, while APLP1 does not contain a KPI domain^31,34^. The KPI-containing soluble form of APP was originally identified as protease nexin-2, and both APP and APLP2 KPI domains have been shown to inhibit several extracellular serine proteases, including trypsin, chymotrypsin, plasmin, kallikreins, and coagulation factor Xia^35–37^. Their protease-inhibitory properties have been demonstrated most clearly for purified KDs and soluble KPI-containing ectodomains, supporting proposed functions in the control of extracellular proteolysis and coagulation.

APP and APLP2 undergo constitutive proteolytic processing by alpha- or beta-secretases, which sheds most of their extracellular region and thereby releases soluble ectodomains that can retain the KPI domain^31,37,38^. Subsequent gamma-secretase cleavage processes the remaining membrane-associated C-terminal fragment^38^. Thus, the APP and APLP2 KDs need not be liberated as isolated peptides to become extracellularly active; they can function as part of larger soluble ectodomains. Whether the KPI domains are also accessible and active while the full-length precursor proteins remain membrane-bound has not been established as clearly. It also remains unknown whether additional proteolytic processing generates free, independently stable APP- or APLP2-derived KDs in vivo.

Importantly, APP and APLP2 are not restricted to the nervous system. APLP2 is broadly expressed, with particularly high transcript abundance reported in human lung, while APP is also expressed in peripheral tissues^33,39,40^. These observations make a role in the respiratory tract biologically plausible, but bulk-tissue expression does not establish which airway cell types produce KPI-containing isoforms, whether the proteins are localized to the apical epithelial surface, or whether infection and inflammation alter their expression, alternative splicing, or ectodomain shedding.

Our findings therefore raise several testable questions: whether KPI-containing APP and APLP2 isoforms are produced by airway epithelial or other respiratory cells; whether full-length surface proteins, shed KPI-containing ectodomains, or smaller proteolytic fragments inhibit TMPRSS2 in vivo; and whether such inhibitors reach effective concentrations in airway surface liquid under homeostatic or inflammatory conditions. Quantifying the relevant isoforms and cleavage products in airway tissue, mucus, and bronchoalveolar fluid, together with testing full-length and shed APP/APLP2 proteins against membrane-associated TMPRSS2, will be necessary to determine whether this activity contributes to endogenous regulation of airway proteolysis and antiviral defense.

Further clinical development of Kunitz domains as host-directed broad-spectrum antivirals is therefore warranted, either in combination with direct-acting antivirals or with agents that limit excessive inflammatory responses. By targeting a conserved host factor required by multiple respiratory viruses, Kunitz domains may impose a higher barrier to viral resistance than direct-acting antivirals alone, while combination therapy could enhance antiviral efficacy, reduce the required doses, and further restrict the emergence of resistant variants.

## Data availability statement

The data that support the findings of this study are available from the corresponding author (J.M.) upon request.

## Ethic statements

Collection of tissue samples for the generation of human airway epithelia cell cultures has been approved by the ethics committee at the Medical School Hannover (airway tissue, application number 2699-2015). All donors provided their informed consent.

## References

1. Laporte, M. & Naesens, L. Airway proteases: an emerging drug target for influenza and other respiratory virus infections. Curr. Opin. Virol. 24, 16–24 (2017).

2. Millet, J. K. & Whittaker, G. R. Host cell proteases: Critical determinants of coronavirus tropism and pathogenesis. Virus Res. 202, 120–134 (2015).

3. Böttcher-Friebertshäuser, E., Klenk, H. D. & Garten, W. Activation of influenza viruses by proteases from host cells and bacteria in the human airway epithelium. Pathog. Dis. 69, 87– 100 (2013).

4. Hoffmann, M. et al. SARS-CoV-2 Cell Entry Depends on ACE2 and TMPRSS2 and Is Blocked by a Clinically Proven Protease Inhibitor. Cell 181, 271–280.e8 (2020).

5. Bertram, S. et al. TMPRSS2 activates the human coronavirus 229E for cathepsin-independent host cell entry and is expressed in viral target cells in the respiratory epithelium. J. Virol. 87, 6150–6160 (2013).

6. Hatesuer, B. et al. Tmprss2 is essential for influenza H1N1 virus pathogenesis in mice. PLoS Pathog. 9, e1003774 (2013).

7. Iwata-Yoshikawa, N. et al. TMPRSS2 Contributes to Virus Spread and Immunopathology in the Airways of Murine Models after Coronavirus Infection. J. Virol. 93, e01815–18 (2019).

8. Keller, C., Böttcher-Friebertshäuser, E. & Lohoff, M. TMPRSS2, a novel host-directed drug target against SARS-CoV-2. Signal Transduct. Target. Ther. 7, 251 (2022).

9. Zhirnov, O. P., Klenk, H. D. & Wright, P. F. Aprotinin and similar protease inhibitors as drugs against influenza. Antiviral Res. 92, 27–36 (2011).

10. Bojkova, D. et al. Aprotinin Inhibits SARS-CoV-2 Replication. Cells 9, 2377 (2020).

11. Wettstein, L. et al. Alpha-1 antitrypsin inhibits TMPRSS2 protease activity and SARS-CoV-2 infection. Nat. Commun. 12, 1–10 (2021).

12. Rawlings, N. D., Tolle, D. P. & Barrett, A. J. Evolutionary families of peptidase inhibitors. Biochemical Journal 378, 705–716 (2004).

13. Laskowski M., Jr. & Kato, I. Protein inhibitors of proteinases. Annu. Rev. Biochem. 49, 593– 626 (1980).

14. Ranasinghe, S. & McManus, D. P. Structure and function of invertebrate Kunitz serine protease inhibitors. Dev. Comp. Immunol. 39, 219–227 (2013).

15. Fries, E. & Blom, A. M. Bikunin--not just a plasma proteinase inhibitor. International Journal of Biochemistry \& Cell Biology 32, 125–137 (2000).

16. Kitaguchi, N., Takahashi, Y., Tokushima, Y., Shiojiri, S. & Ito, H. Novel precursor of Alzheimer’s disease amyloid protein shows protease inhibitory activity. Nature 331, 530–532 (1988).

17. Wasco, W. et al. Isolation and characterization of APLP2 encoding a homologue of the Alzheimer’s associated amyloid beta protein precursor. Nat. Genet. 5, 95–100 (1993).

18. Trexler, M., Bányai, L. & Patthy, L. A human protein containing multiple types of protease-inhibitory modules. Proc. Natl. Acad. Sci. U. S. A. 98, 3705–3709 (2001).

19. Chu, M. L. et al. Mosaic structure of globular domains in the human type VI collagen alpha 3 chain: similarity to von Willebrand factor, fibronectin, actin, salivary proteins and aprotinin type protease inhibitors. EMBO J. 9, 385–393 (1990).

20. de Magalhães, M. T. Q., Mambelli, F. S., Santos, B. P. O., Morais, S. B. & Oliveira, S. C. Serine protease inhibitors containing a Kunitz domain: their role in modulation of host inflammatory responses and parasite survival. Microbes Infect. 20, 606–609 (2018).

21. Lawrenz, J. et al. Trypstatin as a Novel TMPRSS2 Inhibitor with Broad-Spectrum Efficacy against Corona and Influenza Viruses. Advanced Science 12, e2506430 (2025).

22. Lawrenz, J. et al. Severe Acute Respiratory Syndrome Coronavirus 2 Vaccination Boosts Neutralizing Activity Against Seasonal Human Coronaviruses. Clin. Infect. Dis. 75, e653– e661 (2022).

23. Weil, T., Lawrenz, J., Seidel, A., Münch, J. & Müller, J. A. Immunodetection assays for the quantification of seasonal common cold coronaviruses OC43, NL63, or 229E infection confirm nirmatrelvir as broad coronavirus inhibitor. Antiviral Res. 203, 105343 (2022).

24. Reed, L. J. & Muench, H. A simple method of estimating fifty per cent endpoints. Am. J. Epidemiol. 27, 493–497 (1938).

25. Laporte, M. et al. Hemagglutinin Cleavability, Acid Stability, and Temperature Dependence Optimize Influenza B Virus for Replication in Human Airways. J. Virol. 94, (2019).

26. Lawrenz, J. Discovery and characterization of Kunitz inhibitors as antivirals against Corona and Influenza viruses. (Universität Ulm, 2026). doi:10.18725/OPARU-59367.

27. Simmons, G. et al. Inhibitors of cathepsin L prevent severe acute respiratory syndrome coronavirus entry. Proc. Natl. Acad. Sci. U. S. A. 102, 11876–11881 (2005).

28. Kawase, M., Shirato, K., van der Hoek, L., Taguchi, F. & Matsuyama, S. Simultaneous Treatment of Human Bronchial Epithelial Cells with Serine and Cysteine Protease Inhibitors Prevents Severe Acute Respiratory Syndrome Coronavirus Entry. J. Virol. 86, 6537–6545 (2012).

29. Böttcher, E., et al. Proteolytic Activation of Influenza Viruses by Serine Proteases TMPRSS2 and HAT from Human Airway Epithelium. J. Virol. 80, 9896–9898 (2006).

30. Rijsbergen, L. C., van Dijk, L. L. A., Engel, M. F. M., de Vries, R. D. & de Swart, R. L. In Vitro Modelling of Respiratory Virus Infections in Human Airway Epithelial Cells - A Systematic Review. Front. Immunol. 12, 683002 (2021).

31. Zhang, H., Ma, Q., Zhang, Y. & Xu, H. Proteolytic processing of Alzheimer’s β-amyloid precursor protein. J. Neurochem. 120, 9–21 (2012).

32. Jacobsen, K. T. & Iverfeldt, K. Amyloid precursor protein and its homologues: a family of proteolysis-dependent receptors. Cellular and Molecular Life Sciences 66, 2299–2318 (2009).

33. Pandey, P. et al. Amyloid precursor protein and amyloid precursor-like protein 2 in cancer. Oncotarget 7, 19430–19444 (2016).

34. Sandbrink, R., Masters, C. L. & Beyreuther, K. Similar alternative splicing of a non-homologous domain in beta A4-amyloid protein precursor-like proteins. Journal of Biological Chemistry 269, 14227–14234 (1994).

35. Xu, F., Davis, J., Hoos, M. & Van Nostrand, W. E. Mutation of the Kunitz-type proteinase inhibitor domain in the amyloid β-protein precursor abolishes its anti-thrombotic properties in vivo. Thromb. Res. 155, 58–64 (2017).

36. Nostrand, W. E. Van et al. Protease nexin-II, a potent anti-chymotrypsin, shows identity to amyloid β-protein precursor. Nature 341, 546–549 (1989).

37. Oltersdorf, T. et al. The secreted form of the Alzheimer’s amyloid precursor protein with the Kunitz domain is protease nexin-II. Nature 341, 144–147 (1989).

38. Hogl, S., Kuhn, P.-H., Colombo, A. & Lichtenthaler, S. F. Determination of the Proteolytic Cleavage Sites of the Amyloid Precursor-Like Protein 2 by the Proteases ADAM10, BACE1 and γ-Secretase. PLoS One 6, e21337 (2011).

39. Guo, Y., Wang, Q., Chen, S. & Xu, C. Functions of amyloid precursor protein in metabolic diseases. Metabolism 115, 154454 (2021).

40. Sprecher, C. A. et al. Molecular cloning of the cDNA for a human amyloid precursor protein homolog: Evidence for a multigene family. Biochemistry 32, 4481–4486 (1993).

